# Triple Notch/Tgfβ/FoxO1 blockade converts multiple intestinal sub-lineages into β-like cells and lowers glycemia in diabetic animals

**DOI:** 10.1101/2022.03.21.484748

**Authors:** Wen Du, Junqiang Wang, Taiyi Kuo, Liheng Wang, Wendy M. McKimpson, Jinsook Son, Hitoshi Watanabe, Takumi Kitamoto, YunKyoung Lee, Lloyd E. Ratner, Kasi McCune, Ya-Wen Chen, Brendan H. Grubbs, Matthew E. Thornton, Jason Fan, Nishat Sultana, Bryan Diaz, Iyshwarya Balasubramanian, Nan Gao, Sandro Belvedere, Domenico Accili

## Abstract

Insulin is the essential treatment of Type 1 (T1D) and is often used in Type 2 Diabetes. For nearly five decades, efforts have been focused on replenishing β-cells in T1D patients as a more durable treatment. Gut endocrine cells can be converted into insulin-producing cells, but their numbers are limited. In this study we report that insulin-immunoreactive cells with Paneth/goblet cell features are present in human fetal intestine, in addition to enteroendocrine cells. Accordingly, lineage tracing experiments show that, besides enterochromaffin cells, the Paneth/goblet lineage can undergo conversion to the insulin lineage upon genetic or pharmacologic Foxo1 ablation in mice. We leveraged these data to design a screening platform in organoids to accurately quantitate β-like cell reprogramming and fine-tune a combination treatment to increase the efficiency of the conversion process by expanding the intestinal secretory lineage. We identified a triple blockade of FoxO1, Notch, and Tgfβ that, when tested in insulin-deficient diabetic animals resulted in a near-normalization of glucose levels, associated with the appearance of gut insulin-producing cells. The findings illustrate a therapeutic approach to replace insulin treatment in diabetes.

## Introduction

The Center for Disease Control (CDC) estimates that 1.6 million adults and 283,000 children and adolescents have type 1 diabetes in the US alone (https://www.cdc.gov/diabetes/data/statistics-report/diagnosed-diabetes.html). Insulin replacement is a life-saving treatment, but it is not a cure and poses a significant burden on patients and their families. Restoration of endogenous insulin production to cure T1D remains a topic of intense interest (Helman and Melton, 2020). Two alternative approaches have been proposed: transplantation and regeneration.

As early as the 1970s, isolated cadaveric pancreatic islets have been transplanted into T1D recipients to treat diabetes (Najarian et al., 1977). Recent developments in stem cell technologies enabled human embryonic stem cells (hESCs)- or induced pluripotent stem cells (iPSCs)-derived islet replacement (Pagliuca et al., 2014; Rezania et al., 2014; Russ et al., 2015). Mature glucose-responsive β-like cells that are functionally equivalent to cadaveric islets can be obtained by different protocols (Sambathkumar et al., 2018; Sui et al., 2018; Veres et al., 2019). Two studies reporting interim data from ongoing first-in-human iPSCs-based transplants showed restoration of meal-induced C-peptide response for up to 1 year after implantation of iPSCs-derived islet cells, providing proof-of-concept for this approach (Ramzy et al., 2021; Shapiro et al., 2021).

Another potential solution is to convert cell types developmentally related to pancreatic β-cells into functional insulin-secreting cells *in vivo.* However, despite a wealth of reports illustrating conversion of different cell types into β-like cells, poor reproducibility has plagued this area, as well as the intrinsic difficulty of targeting this process pharmacologically (McKimpson and Accili, 2019; Zhou et al., 2008). Although most studies focused on conversion of different pancreatic cell types, there are reports of trans-differentiation from organs developmentally related to the pancreas, such as liver (Banga et al., 2012; Zaret and Grompe, 2008), stomach (Ariyachet et al., 2016), and intestine (Chen et al., 2014).

Genetic ablation of FoxO1 in Neurog3+ progenitor cells can convert enteroendocrine cells (EEC) into insulin-producing β-like cells in mice (Talchai et al., 2012a). Moreover, FOXO1 inhibition using a dominant-negative mutant or lentivirus-encoded small hairpin RNA promotes generation of insulin-secreting cells in human iPSC-derived gut organoids (Bouchi et al., 2014). The potential therapeutic significance of this work was boosted by recent reports: one identifying β-like cells in the human fetal intestine–and thus implying that conversion restores a fetal cell type (Egozi et al., 2021); and the others showing that previously described small molecule FOXO1 inhibitors can yield insulin-producing cells *in vivo* and lower glycemia in diabetic mice (Kitamoto et al., 2022; Lee et al., manuscript in preparation). These findings prompted us to investigate whether other descendants of Neurog3+ progenitors, such as subsets of goblet and Paneth cells also have the potential to be converted into insulin-secreting β-like cells. Based on the identification of cells with mixed lineage of insulin and Paneth/goblet features in human fetal intestines, we developed cellular assays to accurately quantitate cell reprogramming and sought to identify a combination treatment to increase the efficiency of the conversion process by leveraging the expansion of the Neurogenin3 and Paneth/goblet lineages. We found that triple blockade of FoxO1, Notch, and Tgfβ can bolster conversion and result in a robust glucose-lowering effect in streptozotocin diabetic animals. Our findings provide a mechanism underlying intestinal cell trans-differentiation into pancreatic β-like cells and expand its potential therapeutic applications.

## Results

### Intestinal secretory lineage cells express insulin during human fetal development

Pancreas and small intestine share a common endodermal origin. Recent studies show that enteroendocrine K/L cells express insulin during fetal, but not post-natal life (Egozi *et al.*, 2021). These data provide a plausible developmental explanation for the observation that FoxO1 deletion in Neurog3+ endocrine progenitors generates gut β-like, insulin-secreting cells in a cell-autonomous manner (Talchai *et al.*, 2012a), since FoxO1 is generally activated upon terminal differentiation in a variety of cell types (Kitamura et al., 2007; Nakae et al., 2003). We tested the relationship between FoxO1 expression/activity and insulin immunoreactivity in the human fetal intestine. Combined immunohistochemistry and *in situ* hybridization detected cells co-expressing insulin mRNA and protein in fetal human intestine at 15-17 weeks gestational age (GA) (Fig. 1A). Immunostaining also revealed co-reactivity with intestinal secretory cell type markers 5HT (enterochromaffin), lysozyme (Paneth), and GLP-1 (EEC K/L cell) (Fig. 1B). Double-positive 5HT/insulin and lysozyme/insulin cells were detected more frequently at the villi tips in the anterior small intestine, whereas double-positive Glp1/insulin cells were mainly located in the posterior region. No insulin positive cells were found in adult human intestine biopsies (Fig. 1C). Co-staining with FOXO1 indicated that most insulin-positive cells did not express FOXO1, consistent with the possibility that FoxO1 ablation in rodents or organoids recapitulates a developmental stage in human fetal intestine (Fig. 1D, E).

**Figure 1.**
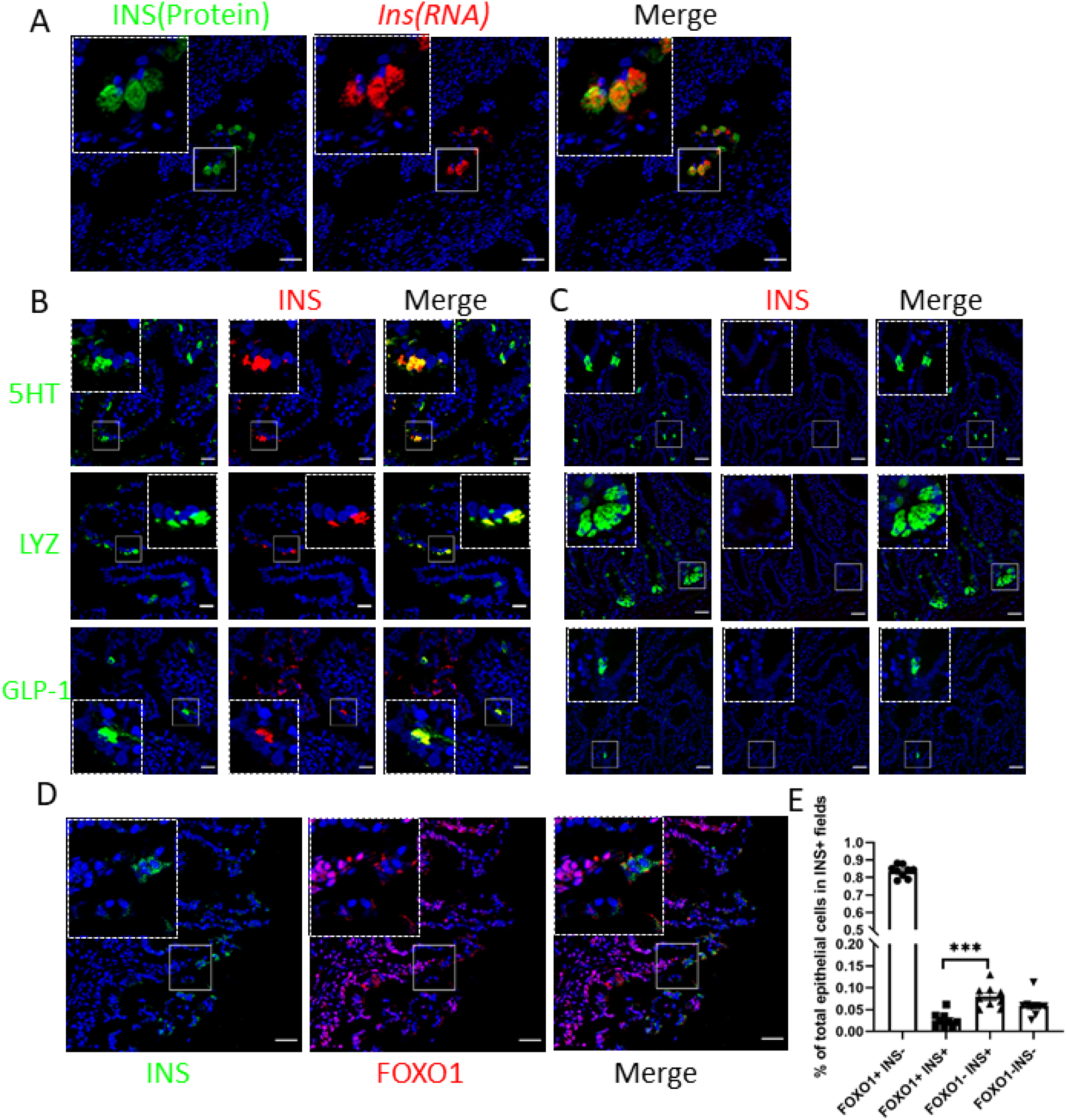
INSULIN and FOXO1 expression in human fetal small intestine secretory lineage cells. (A) (INS-protein, green) and RNAscope (*INS*-mRNA, red) co-staining in fetal human anterior intestine, scale bar = 20μm; (B) Insulin (red) and 5HT, Lysozyme or GLP-1 (green) staining in fetal human anterior intestine, scale bar = 20μm; (C) Insulin (red) and 5HT, Lysozyme or GLP-1 (green) staining in adult human duodenum, scale bar = 40μm; (D) Insulin (green) and FOXO1 (red) staining in fetal human anterior intestine, scale bar = 20μm. (E) Frequency of Foxo1^-^Insulin^+^ vs. Foxo1^+^ Insulin^+^ cells in fetal human anterior intestine (n=10 from 3 different donors) Bar graphs show mean ± SEM t-test, ***, P<0.001).

### Separate subsets of Neurog3 lineage yield intestinal β-like cells

The discovery of cells with mixed insulin/Paneth/goblet features is consistent with the notion that different cell types arise from Neurog3+ progenitors: EEC, goblet, and Paneth cells (Fig. S2A) (Schonhoff et al., 2004). Therefore, we asked whether the latter two subtypes also give rise to gut β-like cells, because this finding would greatly expand the repertoire of target cells for replacement therapy. To answer this question, we used a two-step enrichment procedure of *in vivo* lineage tracing with Neurog3Cre-FoxO1^f/f^; Rosa26^tdtomato^ mice to label FoxO1 knockout cells derived from Neurog3 progenitors (NFKO), followed by CD24 immunostaining to distinguish among EEC, goblet, and Paneth cells (Fig. S1) (Sato et al., 2011b). Quantitative flow cytometry analysis (FACS) revealed that NFKO increased Neurog3-derived cells ~1.7 fold, from 1.54% to 2.65% (P<0.0001) (Fig. 2A, B). Single-cell RNASeq showed an expansion of the EEC and goblet/Paneth lineages among Neurog3 daughter cells of NFKO mice (Fig. S2A, B). Interestingly, both subpopulations included insulin-immunoreactive cells (Fig. 2C). CD24 staining allowed us to subdivide Neurog3-derived (Tomato^+^) cells into two distinct populations: CD24^neg^/Tomato^+^ and CD24^+^/Tomato^+^ (Fig. 2D). CD24^+^ cells included both Paneth and 5HT cells (Fig. S1). QPCR data demonstrated that CD24^neg^/Tomato^+^ cells from NFKO mice were highly enriched in *Ins1* and *Ins2* mRNA (500- to 1,000-fold), while CD24^+^/Neurog3^+^ cells showed a more limited 10- to 30-fold enrichment (Fig. 2E). Consistently, insulin-immunoreactive cells showed weak or absent CD24 membrane staining (Fig. 2F). Notably, Gene Set Enrichment Analysis (GSEA) of pooled RNAseq data showed an increase of pancreatic β-cell- and protein secretion-related transcripts in the CD24^neg^ Tomato^+^ population (Fig. 2G). The morphology and gene expression profiles of these cells was consistent with a dual origin from EEC and Paneth/goblet cells (Fig. 2C, F). Besides Paneth/goblet lineage markers (*Spink4, Defa24, Muc2, Lyz1*) and EEC lineage markers (*Chga, Gcg, Tph1, Cck, Pyy*), we detected quiescent stem cells markers *HOPX* and *Olfm4* in the insulin-immunoreactive population (Fig. S2C, D). Moreover, CytoTRACE analysis showed that the insulin-immunoreactive population represented a less differentiated cells state compared with insulin-negative cells from NFKO mice, supporting the notion that FoxO1 ablation in Neurog3-derived cells brings about a fetal-like stage.

**Figure 2.**
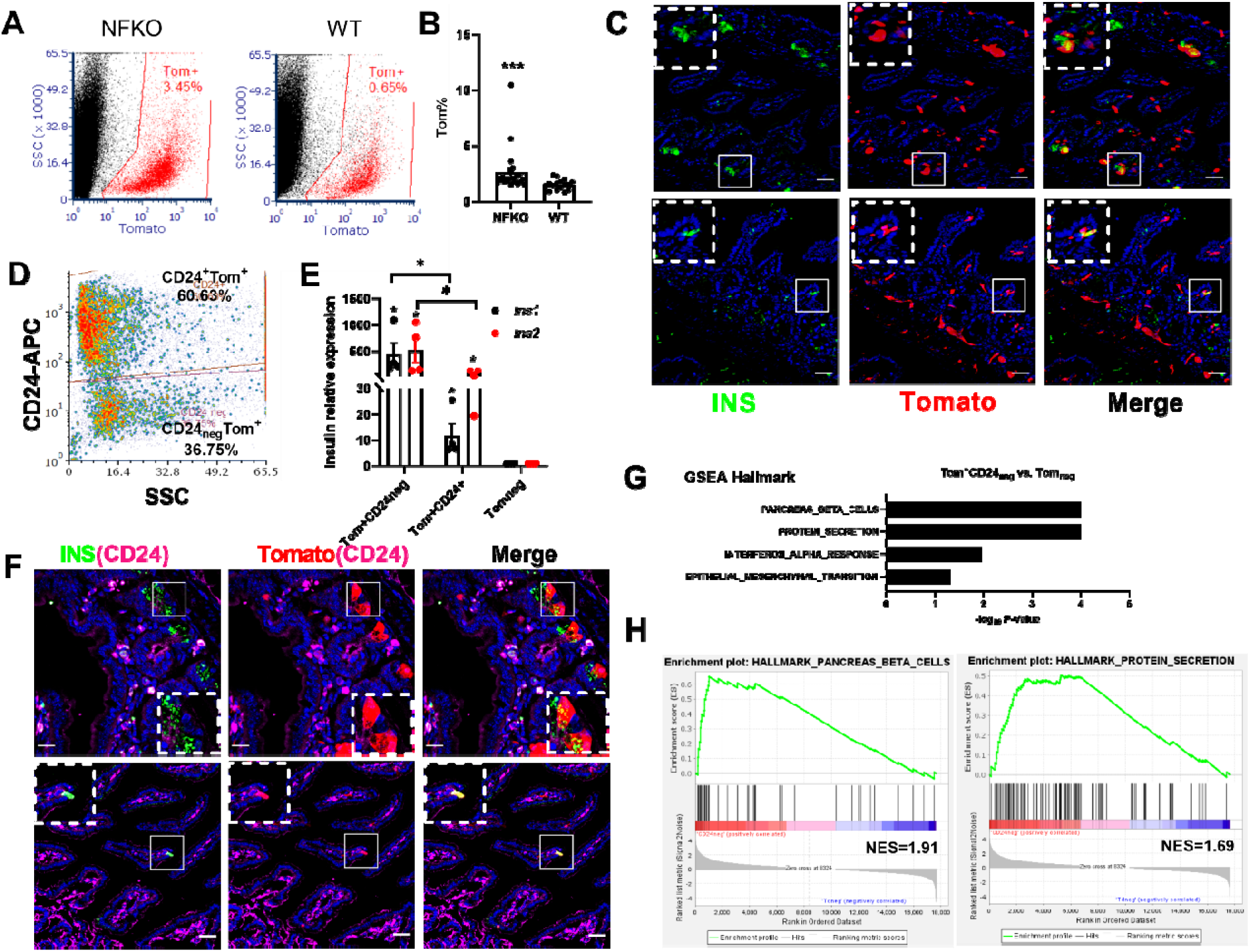
Expanded Neurog3 lineage and β-like cells in gut of Neurog3 FoxO1 KO mice. (A) FACS of isolated Tomato+ cells from either Neurog3Cre-Foxo1^f/f^; ROSA^tdTomato^ (NFKO) or Neurog3Cre+; ROSA^tdTomato^ (Neurog3cre) mouse gut epithelial cells. Red gate indicates sorting window for Neurog3-derived Tomato+ cells; (B) Tomato+ cells frequency assessed by FACS in NFKO and Neurog3Cre (WT) mice (NFKO, n=23; WT, n=16. Bar graphs show mean ± SEM. Student t-test, ***, P<0.001); (C) Representative IHC image of two types of Neurog3-derived β-like cells from NFKO mice: Paneth pattern (upper panel) and EEC pattern (lower panel), scale bar = 40μm; (D) FACS plot CD24 based sorting strategy of dissociated Tomato+ single cells from NFKO small intestinal epithelial cells; (E) *Ins1* and *Ins2* mRNA in sorted CD24^+^Tomato^+^, CD24^neg^ Tomato^+^ and Tomato^neg^ population (n= 4 mice, Mann-Whitney Rank Sum Test, *, P<0.05); (F) Representative IHC of Insulin, CD24 and Tomato. Paneth (upper panel) and EEC pattern (lower panel) of CD24 staining in insulin+ cells; (G-H) Enriched hallmark gene sets in CD24^neg^ Tomato^+^ vs. Tomato^neg^ population predicted by the Gene Set Enrichment Analysis (GSEA).

### Lineage tracing identifies a dual source of β-like cells following FoxO1 ablation

The presence of two distinct cell subtypes with β-like features raised the possibility that other intestinal cell types can be converted to insulin-immunoreactive cells by FoxO1 ablation. To critically test this hypothesis, we assessed generation of β-like cells by tracing the 5HT lineage using primary organoids from Tph1Cre^Ert2/+^; Rosa26^tdTomato^ mice to identify EEC-derived β-like cells, and the goblet/Paneth lineage using Lyz1Cre^Er/+;^ Rosa26^tdTomato^ organoids. After inducing Tph1 reporter-dependent gene activation with 4-hydroxytamoxifen (4-OH-TAM), we enriched organoids in EEC by incubating them in medium containing inhibitors of Notch, WNT, and MEK (DAPT, IWP2, and PD0325901, respectively) (Basak et al., 2017). To induce conversion into β-like cells, we added the chemical FOXO1 inhibitor (AS) (Nagashima et al., 2010; Tanaka et al., 2010). QPCR analysis showed significant increases of *Ins1, Ins2,* and *Tph1* mRNA after incubation in EEC medium with or without AS (Fig. 3A). We found ~11% 5HT cells by immunostaining and flow cytometry (Fig. 3C, D). Treatment with AS increased the percentage of 5HT cells 1.5-fold (Fig. 3C, D). Pulse-chase labelling also showed that after 4-OH-TAM treatment insulin-positive cells colocalized with newly generated 5HT cells (Tomato^+^ cells) regardless of whether organoids had been subjected to the EEC differentiation protocol (Fig. 3D, Fig. S3). Induction of 5HT-positive cells was also evident in EEC-enriched human gut organoids (hGO) (Fig. 3E, F). In sorted 5HT-positive cells from hGOs, insulin mRNA was induced ~ six-fold by incubation in EEC medium, and 30-fold by addition of another FoxO1 inhibitor, FBT10 (Fig. 3G) (Langlet et al., 2017; Lee et al., 2021). These data are consistent with the hypothesis that FOXO1 inhibition facilitates conversion of 5HT cells into β-like cells.

**Figure 3.**
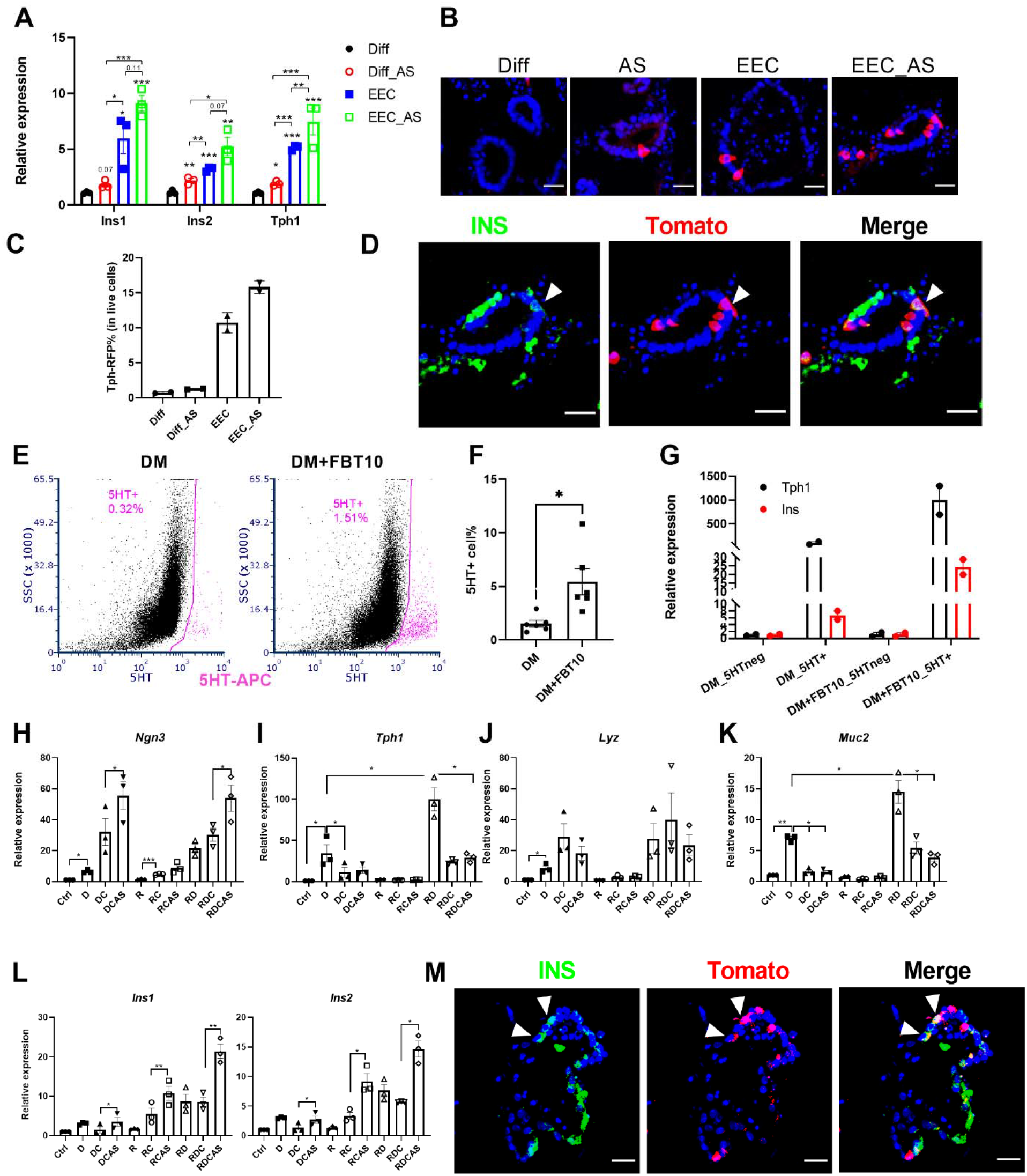
Dual source of β-like cells in murine and human gut organoids. (A) QPCR of mouse intestinal organoids following EEC induction (n=3, bar graphs represent means ± SEM, paired t-test, * P<0.05; ** P<0.01; *** P<0.001); (B) Tomato staining of Tph1Cre^ERT2^ organoids after 4 days in differentiation (Diff) and EEC medium following activation of Tomato reporter (n=3 biological replicates, scale bar: 20 μm). (C) Percentage of Tph1Cre^ERT2^ Tomato cells in mouse gut organoids with or without iFOXO1 (AS1842856, AS) treatment (n=2 biological replicates); (D) Lineage tracing of Tph1Cre^ERT2^ 4 days after activation of Tomato expression (scale bar: 20 μm); (E) FACS diagram representing the sorted 5HT+ (pink) population in EEC induced from human gut organoids with or without FBT10 treatment; (F) Percentage of 5HT+ cells in EEC induced from human gut organoids with or without FBT10 treatment (n=6 biological replicates, bar graphs represent means ± SEM, student t-test, * P<0.05); (G) QPCR of *Ins* and *Tph1* in sorted 5HT+ cells with or without FBT10 treatment (n=2 biological replicates); (H-L) QPCR of different marker genes following treatment with combination of iNotch (DAPT, D), iTGFβ (Repsox, R), iGSK3β (Chir, C); iFOXO1 (AS1842856, AS) (n=3, bar graphs represent means ± SEM, paired t-test, * P<0.05; ** P<0.01; *** P<0.001); (M) Lineage tracing of Lyz1Cre^ER^ 4 days after activation of Tomato expression (scale bar: 20 μm);

Next, we performed similar experiments in organoids derived from Lyz1Cre^Er/+;^ Rosa26^tdTomato^ mice to label goblet/Paneth cells, followed by induction of these two interrelated lineages (Fig. 3H). We optimized chemical induction of the Paneth/goblet lineage by different combinations of the GSK3β inhibitor Chir99021, Notch inhibitor DAPT, and TGFβ inhibitor Repsox. QPCR data showed that single Notch inhibition enriched all secretory cell markers, such as *Neurog3, Tph1, Lyz1* and *Muc2;* this effect was strengthened by the TGFβ inhibitor. In contrast, addition of the GSK3β inhibitor increased *Neurog3* and *Lyz1* but decreased *Tph1* and *Muc2,* consistent with the possibility that this combination promotes an earlier stage of EEC differentiation. Single treatment with either TGFβ or GSK3 inhibitor had no effect (Fig. 3H-K). The triple combination of Notch, TGFβ, and GSK3β inhibitors resulted in a 10-fold induction of *Ins1* and *Ins2* mRNA along with EEC progenitor markers, indicating a trend toward the β-cell-like phenotype. The effect of the triple blockade was amplified by adding the FOXO1 inhibitor AS, with both *Ins1* and *Ins2* mRNA expression increasing by 20- and 14-fold, respectively (Fig. 3L). Adding AS also strengthen the effect of the dual Notch/TGFβ or Notch/GSK3β blockade (Fig. 3L). To confirm the origin of β-like cells, we performed immunohistochemistry and found that insulin colocalized with LYZ1-tomato cells, indicating that β-like cells can also arise from Paneth/goblet cells (Fig. 3M).

### Newly developed FOXO1 inhibitors increase β-like cell conversion

Based on these data, we sought to increase the efficiency of generating β-like cells using a modified secretory cell conversion protocol combined with FoxO1 inhibition in mouse intestinal organoids (Jeffrey Michael KARP, 2017) (Fig. 4A). TGFβ inhibition resulted in strong induction of *Ins1* and *Ins2;* the latter was further increased ~ twofold by the FoxO1 inhibitor, AS. Notch inhibition had a stronger effect on *Ins1* than on *Ins2,* and addition of AS increased both. Addition of the Notch inhibitor to the TGFβ blockade, either in the presence or absence of AS, had no effect, indicating that Notch is epistatic to TGFβ in the pathway leading to β-like cell conversion (Fig. 4B, Fig. S4). We interpret these data as suggesting that Notch signaling affects primarily β-cell differentiation, while TGFβ signaling affects β-cell maturation. Measurements of insulin content in organoid extracts are consistent with the mRNA findings and show that FOXO1 inhibition increased the amount of insulin recovered after single TGFβ or dual Notch/TGFβ blockade (Fig. 4C).

**Figure 4.**
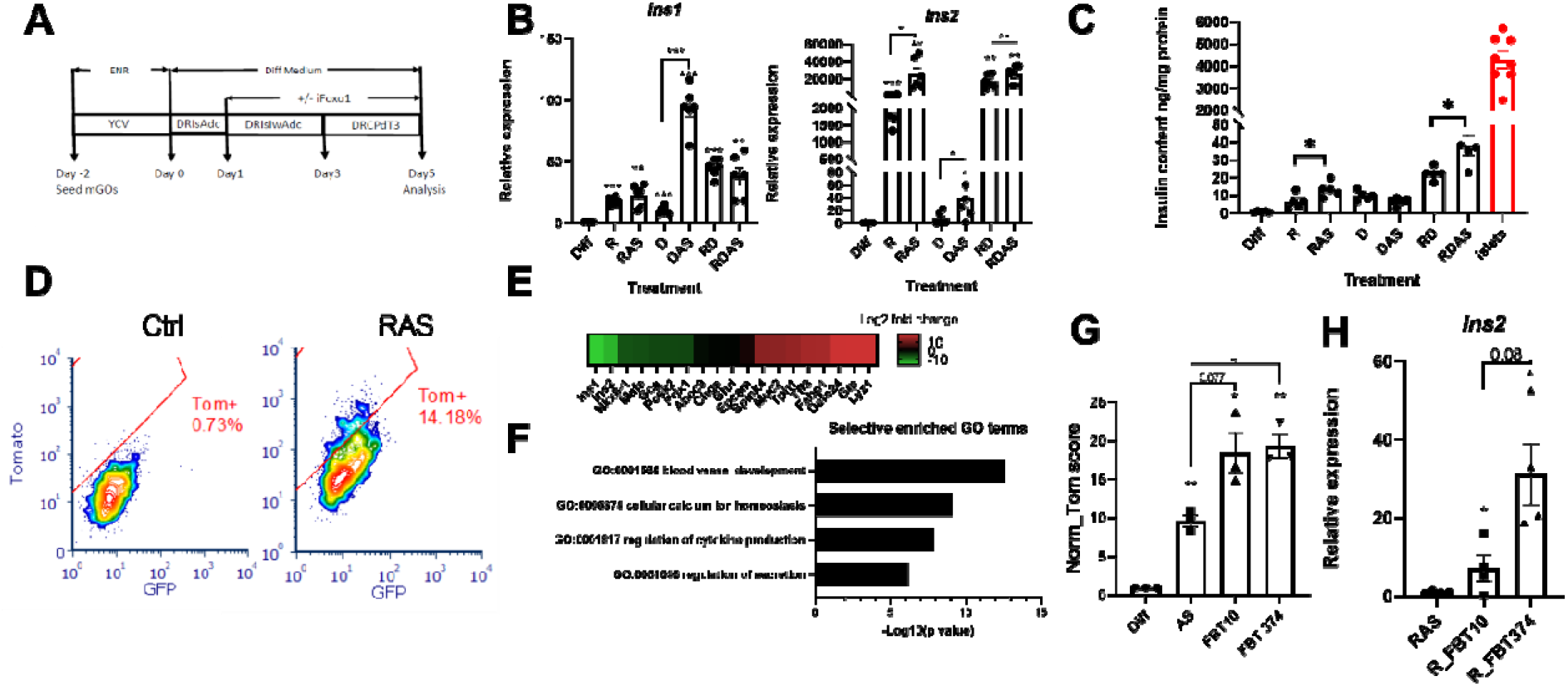
Conversion of gut cells into insulin-producing cells by combination treatment. (A) Modified differentiation protocol to induce conversion of INS+ cells by the addition of FOXO1 inhibitor to the differentiation medium at different stages; (B) QPCR of *Ins1 and Ins2* expression from organoids following treatment with the differentiation cocktail, comprising a combination of WNT, TGFβ, and DNA methyltransferase inhibitors, along with chemical inducers of Neurogenin3, followed by the addition of FOXO1 inhibitor (n=6, bar graphs represent means ±SEM, paired t-test, * P<0.05; ** P<0.01; *** P<0.001); (C) Insulin content in organoids following treatment with the differentiation cocktail compared with islets (n=4, biological replicates, bar graphs represent means ± SEM, one-way ANOVA, * P<0.05); (D) FACS diagram presenting the percentage of converted INS2+ cells in ctrl vs. differentiation cocktail-treated organoids; (E) Heatmap comparing expression level of typical islet and gut epithelial marker genes in sorted Tom^+^ cells from islets vs. differentiated organoids of INS2-Tomato mice; (F) GO analysis of upregulated pathways in sorted Tom+ cells from islets vs. differentiated organoids of INS2-Tomato mice; (G) Comparative evaluation of the potencies of 2 new FBT compounds and iFOXO1 (AS1842856) to generate Insulin+ cells by an integrated calculation (Tom Score) of INS2-Tomato intensity, percentage of INS2-Tomato cells, and live-cell percentage, as detected by flow cytometry (n=3 biological replicates, bar graphs represent means ± SEM, paired t-test, * P<0.05; ** P<0.01); (H) *Ins2* relative expression in sorted INS2-Tomato cells (n=4, bar graphs represent means ± SEM, paired t-test, * P<0.05).

Next, we set up quantitative assays with cultured gut organoids derived from mice bearing a Rip-Cre; Rosa26^tdtomato^ reporter allele to evaluate the efficiency of β-like cell conversion. In this assay, we used FACS to separate and quantitate cells in which *Ins2* expression had been activated, as indicated by the Tomato reporter, from non-insulin-expressing epithelial cells. In a typical experiment, about 14% of cultured organoid cells demonstrated onset of red fluorescence following combined TGFβ/FoxO1 inhibition (Fig. 4D). Using this screening platform, we validated several new novel FoxO1 inhibitors (FBT) based on their potency in reporter promoter assays (Lee et al., manuscript in preparation). We selected two compounds, FBT10 and FBT374, that outperformed AS with regard to conversion frequency to β-like cells and *Ins2* expression (Fig. 4G,H). Using Tomato-positive (β-like) cells isolated by FACS, we compared expression of β-cell-specific genes between converted β-like cells and islet β-cells. RNAseq showed that organoid-derived insulin-immunoreactive cells express pancreatic β-cell markers, including *Nkx6.1, MafA, Pdx1,* and *Pcsk2,* while also retaining some intestinal epithelial markers (Fig. 4E). Gene ontology (GO) analysis revealed that differentially expressed genes (DEGs) were related to biological processes in pancreatic β-cells including calcium homeostasis and hormone secretion (Fig. 4F). Thus, intestinal β-like-cells obtained by FOXO1 inhibition share a common molecular signature with islet β-cells.

### Inhibition of Notch and TGFβ in FoxO1-deficient mice increases Neurog3+ and β-like cells

Next, we tested whether triple blockade of Notch, TGFβ and FOXO1 can induce cell conversion *in vivo.* To this end, we combined genetic FoxO1 ablation in Neurog3cre-FoxO1^f/f^; Rosa26^tdtomato^ mice with pharmacological treatment with Repsox and a different Notch inhibitor, the γ-secretase inhibitor, Dibenzazepine (DBZ). We injected DBZ for the first 2 days, followed by 5 days of Repsox oral dosing (Fig. 5A). Immunohistochemistry and quantitative FACS showed that DBZ increased Neurog3-derived cells by ~ 2.5-fold, and the DBZ/Repsox combination by 7-fold, to account for ~ 15% of all live duodenal epithelial cells. Repsox had no effect by itself (Fig. 5C-D). The treatment had similar effects on the percentage of 5HT cells, consistent with data in cultured organoids (Fig. 5E-F). These data show that triple inhibition expanded the EEC lineage, a necessary condition for β-like conversion of a subset of cells.

**Figure 5.**
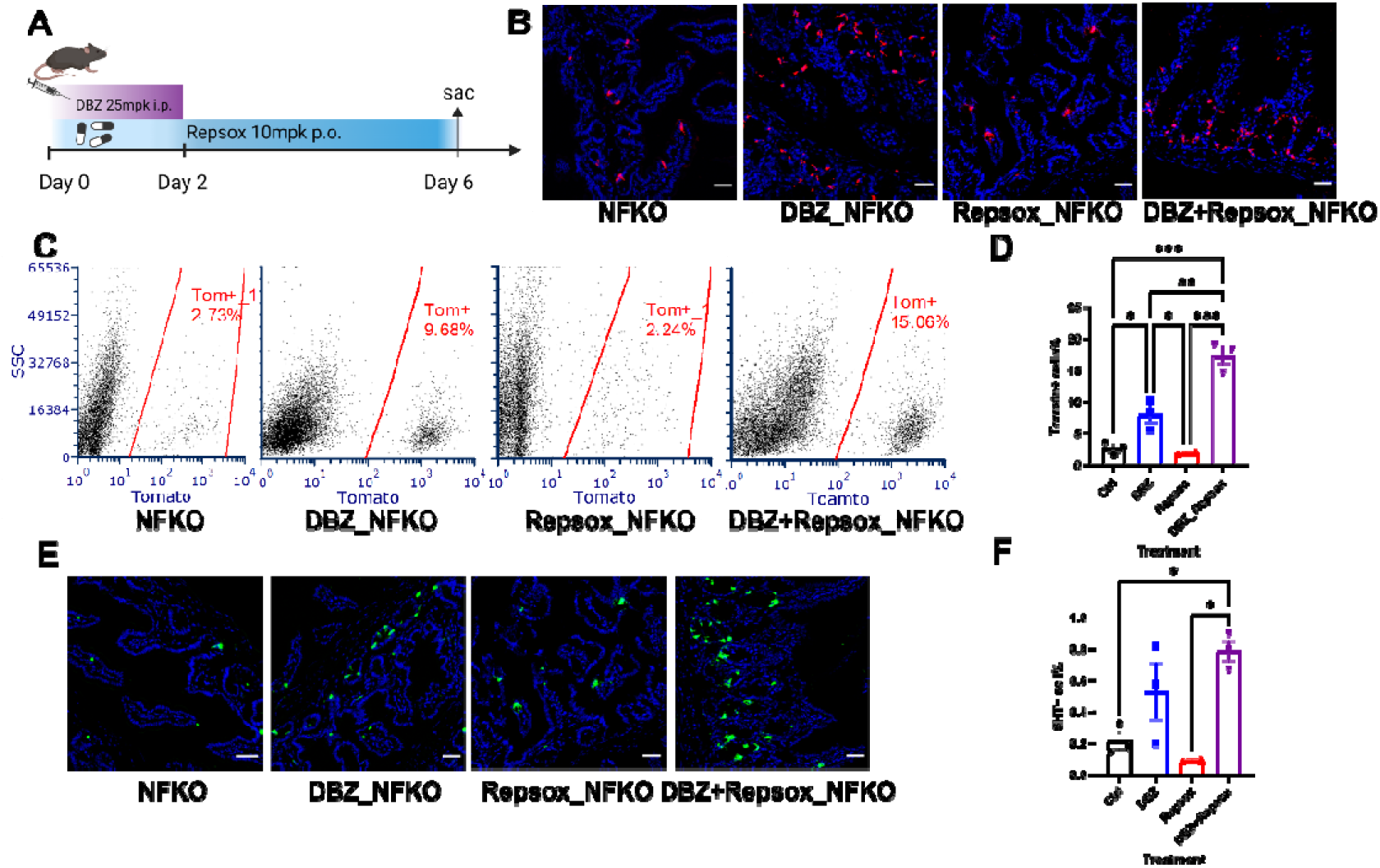
Notch and TGFβ inhibitors expand the Neurog3 lineage in NFKO mice. (A) Experimental design for DBZ and Repsox combination treatment of NFKO mice; (B) Representative IHC image of Tomato (Red) staining in Neurog3Cre FoxO1^f/f^;ROSA^tdTomato^ mice following DBZ, Repsox or combination treatment, scale bar: 40uM; (C) Representative flow cytometry of Tomato+ cells sorting from Neurog3Cre FoxO1^f/f^;ROSA^tdTomato^ mice following DBZ, Repsox or combination treatment; (D) Tomato+ cells frequency measured by FACS in Ctrl, DBZ-, Repsox- or combination-treated Neurog3Cre Foxo1^f/f^;ROSA^tdTomato^ mice; (E) Representative IHC image of 5HT (green) staining in duodenum from Neurog3Cre FoxO1^f/f^;ROSA^tdTomato^ following treatment with DBZ, Repsox or combination, scale bar: 40uM; (F) Quantitation of 5HT+ cells by intracellular staining of FACS sorts in Ctrl, DBZ-, Repsox- or combination-Neurog3Cre FoxO1^f/f^;ROSA^tdTomato^ mice; n=3 mice in each treatment group. Bar graphs represent means ± SEM. * P<0.05; ** P<0.01; *** P<0.001 by two-way ANOVA.

To test whether these treatments resulted in the formation of functional gut β-like cells, we rendered the animals diabetic through ablation of pancreatic β-cells by STZ and measured the effects of the various interventions on fasting glucose, glucose tolerance, plasma insulin, and generation of insulin-immunoreactive intestinal cells (Fig. 6A). FoxO1 ablation resulted in lower fasting glycemia and improved GTT after STZ administration, consistent with prior observations (Talchai *et al.*, 2012a). After a 6-day course, adding dual inhibition on top of the FoxO1 knockout increased plasma insulin levels in 4-hr-fasted or 1-hr-refed mice (Fig. 6B, C). In oral glucose tolerance tests, dual inhibition in NFKO mice yielded the largest improvement compared with vehicle and any single treatment, including a trend with dual inhibition in WT controls (Fig. 6D, E). Immunohistochemistry revealed abundant insulin-immunoreactive cells within the intestines located mainly in crypts, and colocalizing with 5HT and lysozyme/MUC2 (Fig. 6F). These data indicate that combined FOXO1, Notch, and TGFβ inhibition increases the efficiency of cell conversion *in vivo* and is associated with a commensurate glucose-lowering effect in diabetic animals.

**Figure 6.**
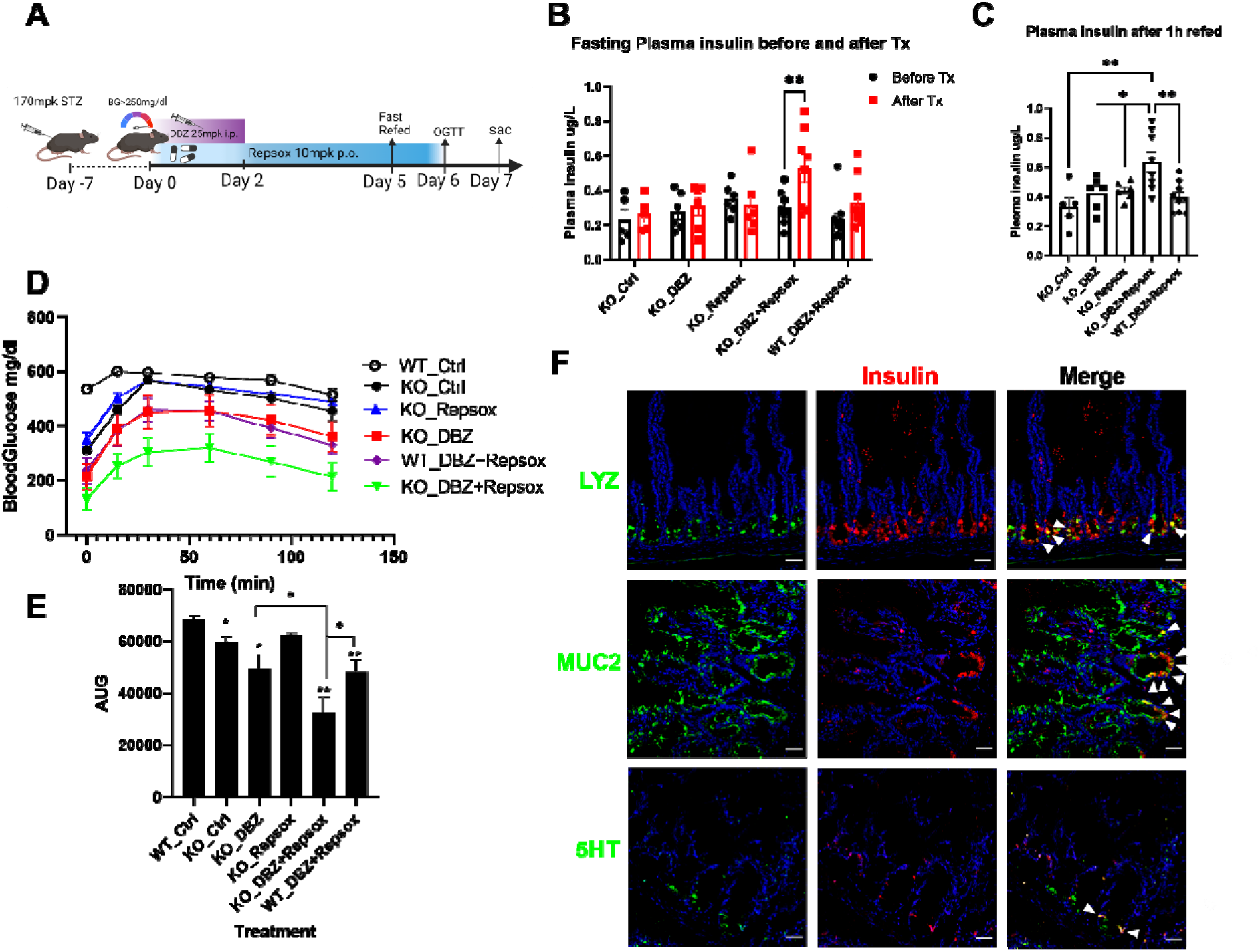
iNotch and iTGFβ combination therapy generates insulin+ cells in NFKO mice. (A) Experimental design for DBZ and Repsox combination treatment in STZ-WT or NFKO mice; (B) 4-hour fasting plasma insulin levels in STZ-NFKO and STZ-WT mice before and after treatment with the indicated compounds; (C) 1-hour refed plasma insulin levels in STZ-NFKO and STZ-WT mice before and after treatment with the indicated compounds; (D) Intraperitoneal glucose tolerance tests after DBZ, Repsox and dual treatment; (E) Area under the curve (AUC) of intraperitoneal glucose tolerance test shown in (E); (F) Representative IHC image of lysozyme (upper panel, green), MUC2 (middle panel, green) and 5HT (lower panel, green) co-stained with Insulin (red) in combination therapy-treated STZ-NFKO mice, scale bar=40μm. N = 6 to 9 mice in each treatment group, bar graphs indicate mean± SEM, two way ANOVA, * P<0.05; ** P<0.01; *** P<0.001.

### Combination treatment of human gut organoids induces insulin+ cells

We finally determined the effects of triple chemical blockade of Notch, TGFβ and FOXO1 using primary human duodenal organoids (Fig. 7A). QPCR analysis indicated a remarkable induction of insulin and CD49a, a membrane marker of human stem cell-derived β-cells by the triple combination treatment (Fig. 7B). Measurements of insulin content and C-peptide immunohistochemistry confirmed these findings (Fig. 7C, D). Thus, triple blockade can robustly induce β-like cell conversion in human organoids.

**Figure 7.**
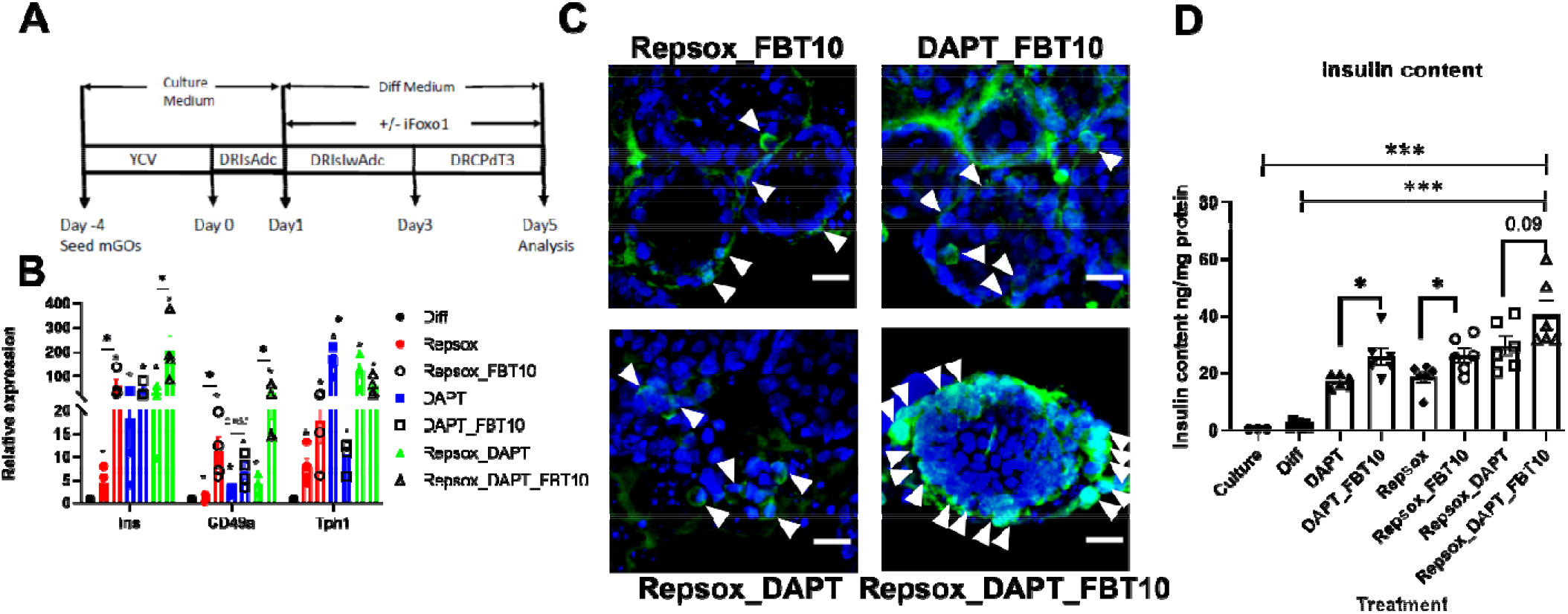
Induction of β-like cells by combination treatment in human gut organoids. (A) Schematic treatment protocol; (B) qPCR of different marker genes from hGOs treated with the differentiation cocktail, bar graphs represent means ± SEM, paired t-test, * P<0.05; (C) Representative IHC image of C-peptide (green) staining in hGOs treated with differentiation cocktail, scale bar: 20 μm; (D) Insulin content of the differentiation cocktail treated hGO (n=6 biological replicates, bar graphs represent means ± SEM, one-way ANOVA, * P<0.05, ** P<0.01, *** P<0.001).

## Discussion

Pancreas and intestine share a common developmental origin, and their endocrine compartments share a common progenitor as well as several terminally differentiated cell types, such as SST- and Ghrelin-producing cells. Other cells, for example α- and K/L cells, give rise to alternatively spliced products of the same pre-proglucagon gene in the two organs (Holst, 2019). Insulin-producing cells are however restricted to the pancreas. We were therefore quite surprised when a decade ago we observed that genetic ablation of FoxO1 in endocrine progenitors resulted in the generation of intestinal cells with highly differentiated β-like cell features as they are only found in pancreatic islets. Three subsequent pieces of evidence clarified this finding. First, Stanger and Zhou independently replicated these findings using a forced expression approach with Neurog3, Pdx1, and MafA (Ariyachet et al., 2016; Chen et al., 2014), confirming the potential of the gut to undergo this conversion. Next, we showed that FOXO1 inhibition appeared to reprogram enterochromaffin 5HT cells into β-like cells in hGO (Bouchi *et al.*, 2014). And the Melton laboratory reported that enterochromaffin-like cells are a “byproduct” of stem cell differentiation into β-cells, including expression of genes related to serotonin biosynthesis. The similarities between these two cell populations suggest that there is a relationship between enterochromaffin and β-cell fates (Veres *et al.*, 2019). This, in turn, dovetails with the notion that pancreatic β-cells synthesize 5HT (Kim et al., 2010). Finally, the recent description of *bona fide* insulin-producing cells in the fetal human gut suggests that FOXO1 ablation is arresting Neurog3 progenitor cell differentiation at a fetal-like stage (Egozi *et al.*, 2021), providing a plausible underpinning for these disparate observations.

One unexplained feature of these findings was that other secretory lineage cells also arise from Neurog3 progenitors (Schonhoff *et al.*, 2004). In this regard, it is noteworthy that the non-canonical Wnt/planar cell polarity pathway, which controls islet β-cell functional heterogeneity, primes intestinal stem cells toward the EEC or Paneth lineages (Böttcher et al., 2021). In this work, we provide direct lineage tracing evidence that secretory cells in the non-endocrine (Paneth and goblet) fate can also be converted to intestinal β-like cells. In addition, expanding on Egozi (Egozi *et al.*, 2021), we show that in human fetal intestine of 15- to 17-week gestational age, insulin-immunoreactive cells also colocalize with goblet/Paneth lineage markers but exclude active FOXO1, lending further support to the notion that FOXO1-inactive cells can be converted to β-like cells. These findings address the question of which type of cell can be converted into insulin-immunoreactive β-like cells, extending previous observations (Bouchi *et al.*, 2014; Kitamoto et al., 2021; Talchai *et al.*, 2012a).

The data above raised the possibility that leveraging additional signaling pathways can modulate the conversion process in synergy with FOXO1. Transforming Growth Factor-β (TGF-β), Wnt, fibroblast growth factor (FGF), Notch, bone morphogenic protein (BMP), and FoxO1, along with relevant receptors and signaling pathways, are involved in pancreatic and intestinal tissue patterning (Boonekamp et al., 2020; Nostro et al., 2011). FoxO1 and Notch signaling interact in determining intestinal stem cell differentiation into Paneth/goblet (Ludikhuize et al., 2020) and EEC lineages (Kitamoto *et al.*, 2021; Zeve et al., 2022). Thus, we combined genetic FoxO1 knockout with pharmacological Notch inhibition (DBZ) to show that dual Notch/FoxO1 inhibition expands the Neurog3+ progenitor pool and its secretory lineage cell descendants. Moreover, adding the TGFβ inhibitor Repsox further increased the Neurog3+ lineage, indicating a synergistic effect on endocrine induction, similar to observations in protocols to derive β-like cells from stem cells (Rezania *et al.*, 2014). Interestingly, TGFβ inhibition in the combination treatment decreased expression of the Paneth cell marker lysozyme, but increases the EC marker, 5HT, and the goblet marker, MUC2, indicating that the two pathways affect sub-lineage specification.

Numerous studies tracking the fate of Paneth, goblet, tuft, and enteroendocrine cells have shown that lineage-committed cells are capable of dedifferentiating into multipotent ISCs during gut regeneration (Beumer et al., 2020; de Sousa and de Sauvage, 2019; Liu and Chen, 2020). Dedifferentiated cells can adopt an alternate cell fate upon injury or perturbation of the intestinal epithelium. The molecular mechanisms driving cellular reprogramming remain to be elucidated. Our findings strengthen the notion that FOXO1 participates in intestinal secretory lineage trans-differentiation, similar to its role in pancreatic β-cells (Son et al., 2021; Talchai et al., 2012b). Single cell RNA-seq of FoxO1-ablated cells also shows the reemergence of HOXP- and Olfm4-positive cells along with β-like cells, suggesting that committed secretory cells can revert to a stem- or fetal-like stage as a path to differentiate into β-like cells. This process too bears similarities with the role of FoxO1 in pancreatic islets (Talchai *et al.*, 2012a).

### LIMITATIONS OF STUDY

Combination treatment had a major glucose-lowering effect in diabetic mice, adding to an emerging body of evidence that pharmacological FoxO1 inhibition is a viable option for β-cell replacement. Although our focus was to probe the mechanistic underpinning of the conversion, the potential use of combination treatment as an alternative to insulin injections or cell transplant should be considered. Most modern treatments leverage detailed knowledge of signaling pathways to target disease processes as diverse as different types of cancer of immune disorders. The advantages of a β-cell replacement that doesn’t entail wearing devices to infuse insulin and measure glucose, or multiple daily insulin injections, with attendant risk of hypoglycemia, can potentially offset regulatory approval hurdles.

In summary, we characterized insulin-secreting β-like cells using genetic and pharmacologic models of signaling perturbations. Based on this insight, we developed a robust combination treatment to generate β-like gut cells in mice and cultured human enteroids. The discovery of similar cells in the human fetal intestine (Egozi *et al.*, 2021) raises the question of whether these manipulations restore a fetal-like cell type. In addition to providing developmental and mechanistic insight into this process, our findings expand potential therapeutic options for insulin replacement in diabetes.

## Acknowledgements

We thank members of the Accili laboratory for helpful discussions and critical data review and T. Kolar and A.M. Flete for technical help. We thank Forkhead BioTherapeutics Inc for Providing FBT compounds. We thank Drs. Que (Columbia University) and Merchant (University of Arizona) for donating mice, Kuo (Stanford University) and Clevers (Hubrecht Institute) for cell lines. We thank M. L. Wilson (Department of Preventive Medicine, University of Southern California and Family Planning Associates) for coordinating fetal tissue collection. Mrs. Xu and Sun performed histology, Drs. Lu and Chen flow cytometry. We thank Dr. Leibel for insightful discussions of the data. This research was supported by a grant from the JPB Foundation to D.A., R01DK119198 to N.G., and by Core Facilities supported by DK63608 (Columbia Diabetes Center).

## Author contributions

W.D. designed, executed experiments, analyzed results, and wrote the manuscript; J.W. analyzed the sc-Seq data; T.K., W.L., W.M.M., J.S., H.W., T.K., L.Y., N.S., B.D. performed experiments and edited the manuscript; Y.C., B.H.G and M.E.T collected the fetal tissue and perform the human fetal tissue related experiment; L.E.R. and K.M. performed surgeries from which donor samples were obtained; N.G. and L.B. maintain the mouse stain; L.Y. and S.B. provide FBT compounds in this study; D.A. designed experiments, oversaw research and wrote the manuscript.

---

**Figure S1.**
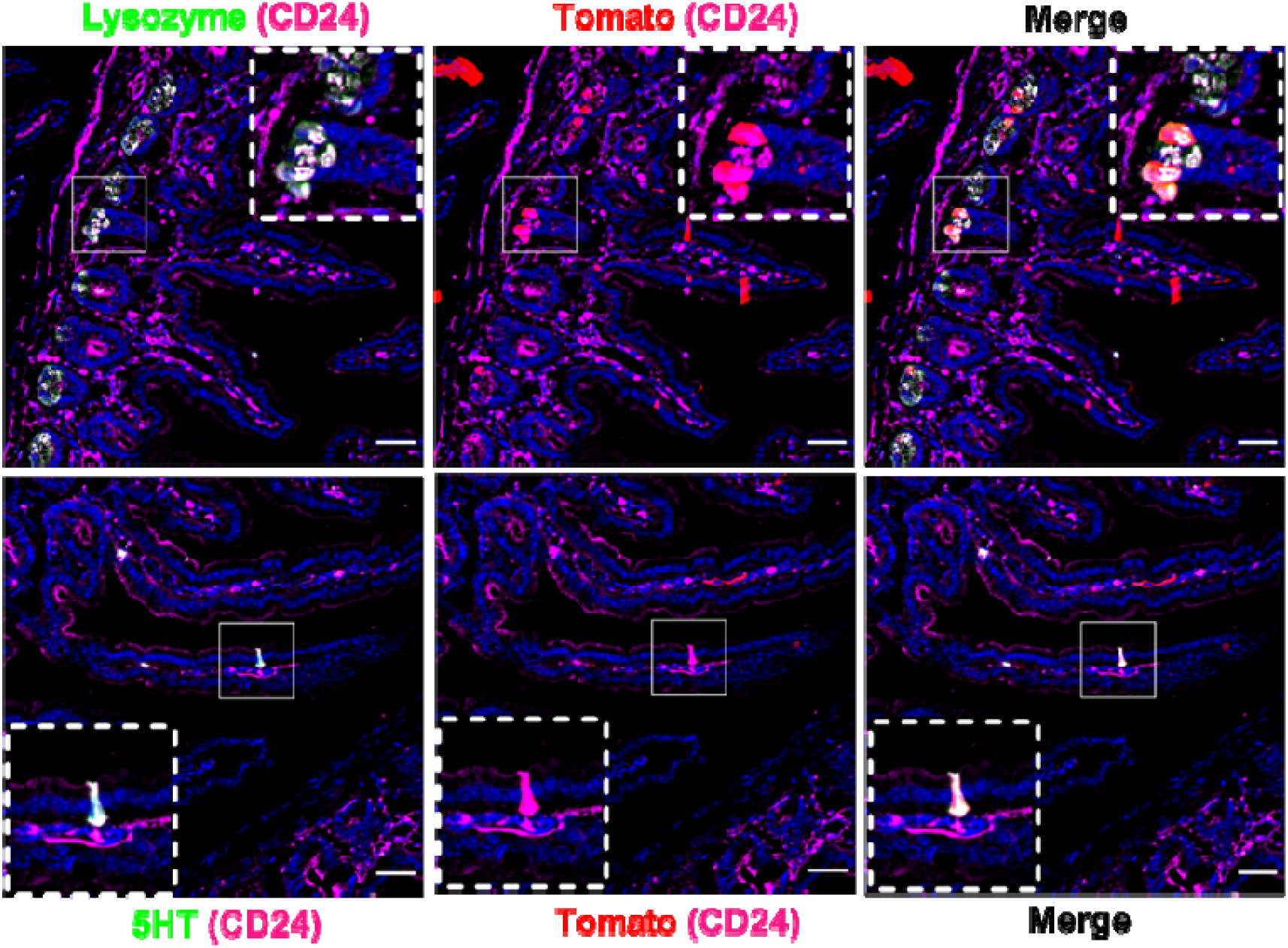
Neurog3-derived canonical 5HT- and lysozyme-immunoreactive cells are CD24^+^ Immunohistochemistry of Tomato, CD24, Lysozyme (upper panel), or 5HT (lower panel). Lysozyme or 5HT (green), Tomato (red), CD24 (magenta), scale bar: 40uM, n=3 mice.

**Figure S2.**
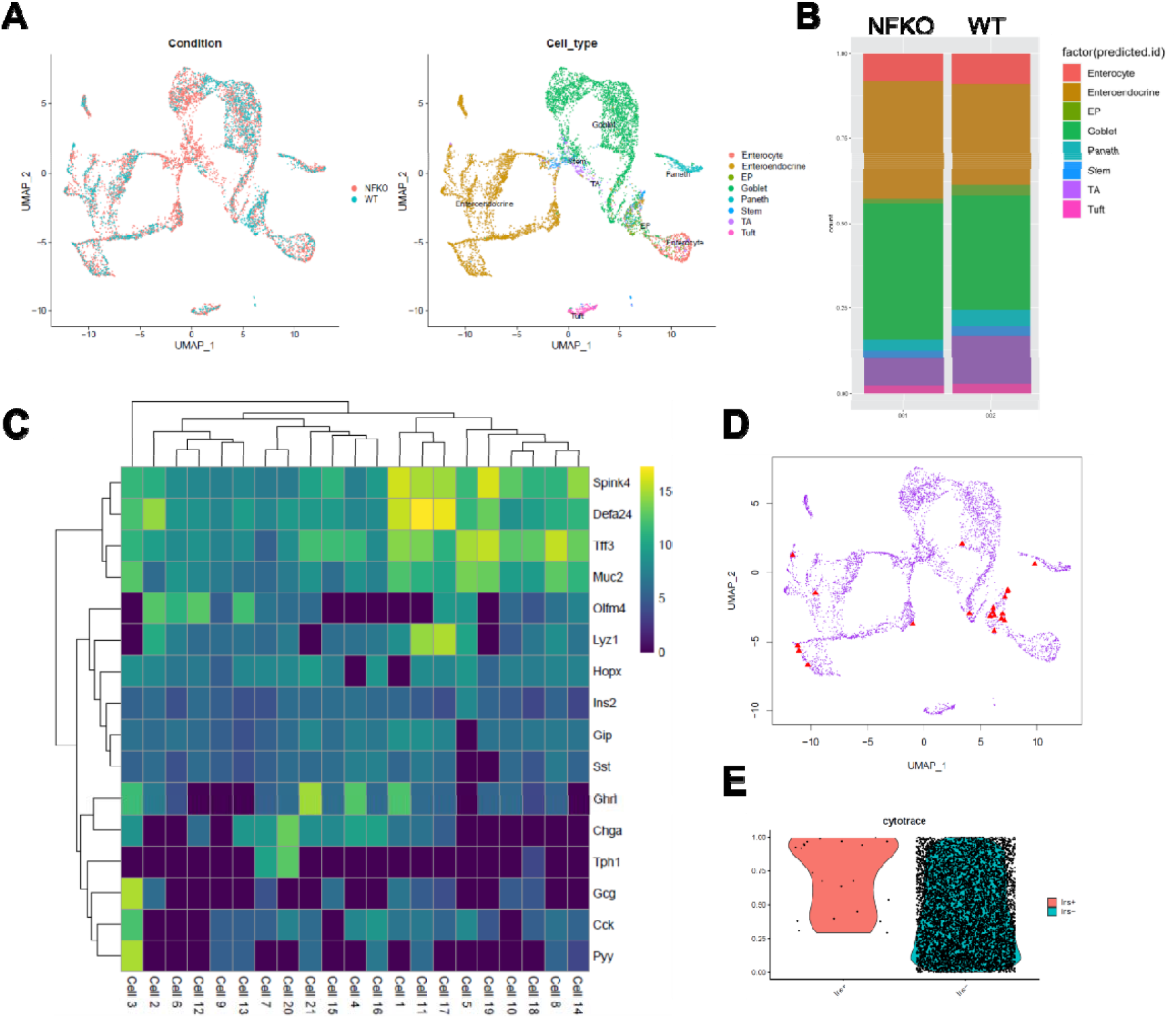
Expanded EEC and goblet-Paneth lineages in NFKO by scRNAseq of Neurog3-derived cells. (A) UMAP plots comparing of NFKO and WT cells clustering (Left, sorted from 4 mice) and predicted cell type identities (Right); (B) Quantification of cell-type composition in NFKO and WT Neurog3-Tomato cells; (C) Marker gene expression heatmap and hierarchical clustering of Insulin+ cells in sorted NFKO-Tomato cells; (D) UMAP plots of Neurog3-Tomato cells from NFKO mice popup of INS expression cells. The Insulin^+^ cells are marked by red triangles; (E) Violin plot of predicted stemness scores in INS^+^ cell group and INS^-^ cell group. Each cell is visualized as a point. The stemness scores are inferred by CytoTRACE.

**Figure S3.**
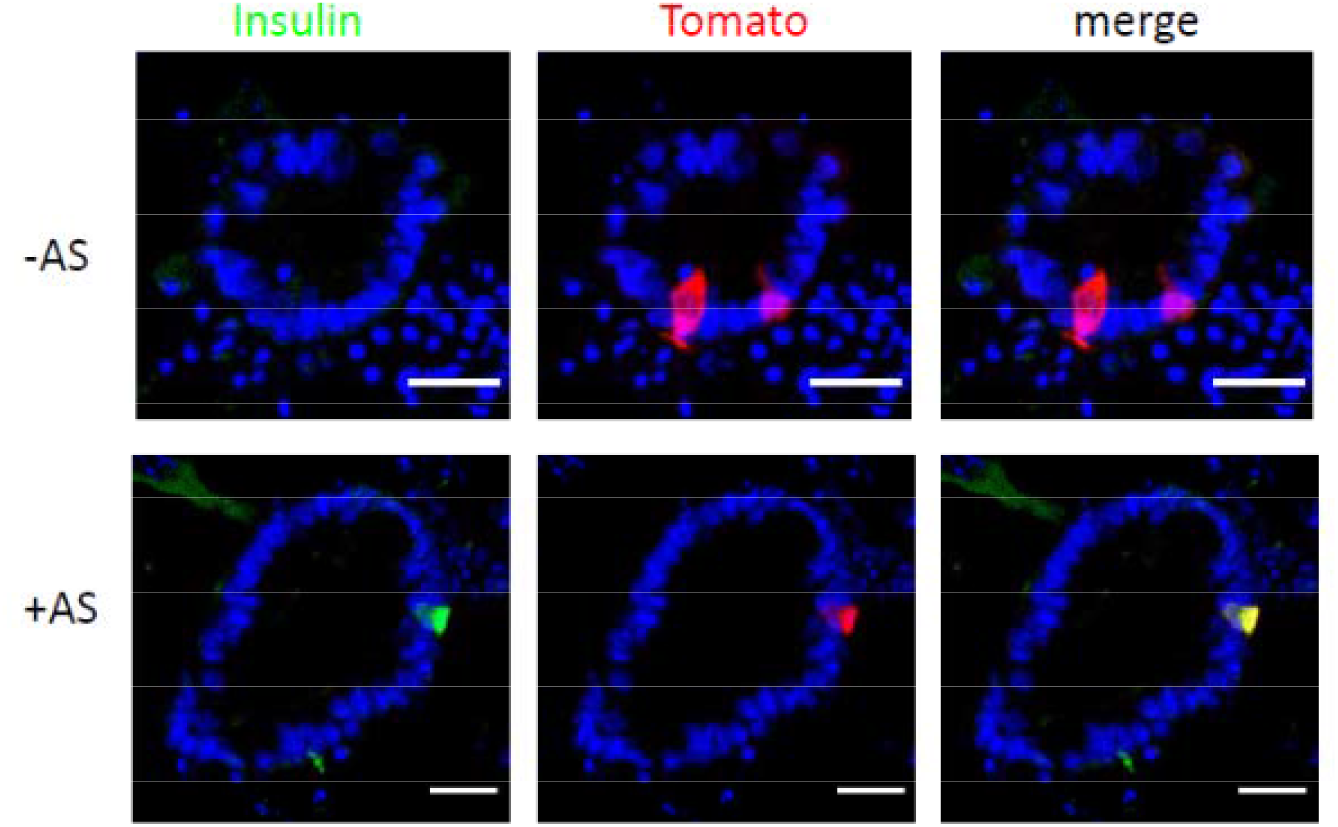
iFOXO1 generates 5HT cell-derived insulin+ cells independent of Notch inhibition. Insulin and Tomato double staining of lineage-traced organoids from Tph1Cre^ERT2^ after four days incubation in differentiation (Diff) medium following induction of Tomato reporter with (lower panel) or without (upper panel) iFoxO1 (AS) treatment (n=3, biological replicates, scale bar: 20 μm).

**Figure S4.**
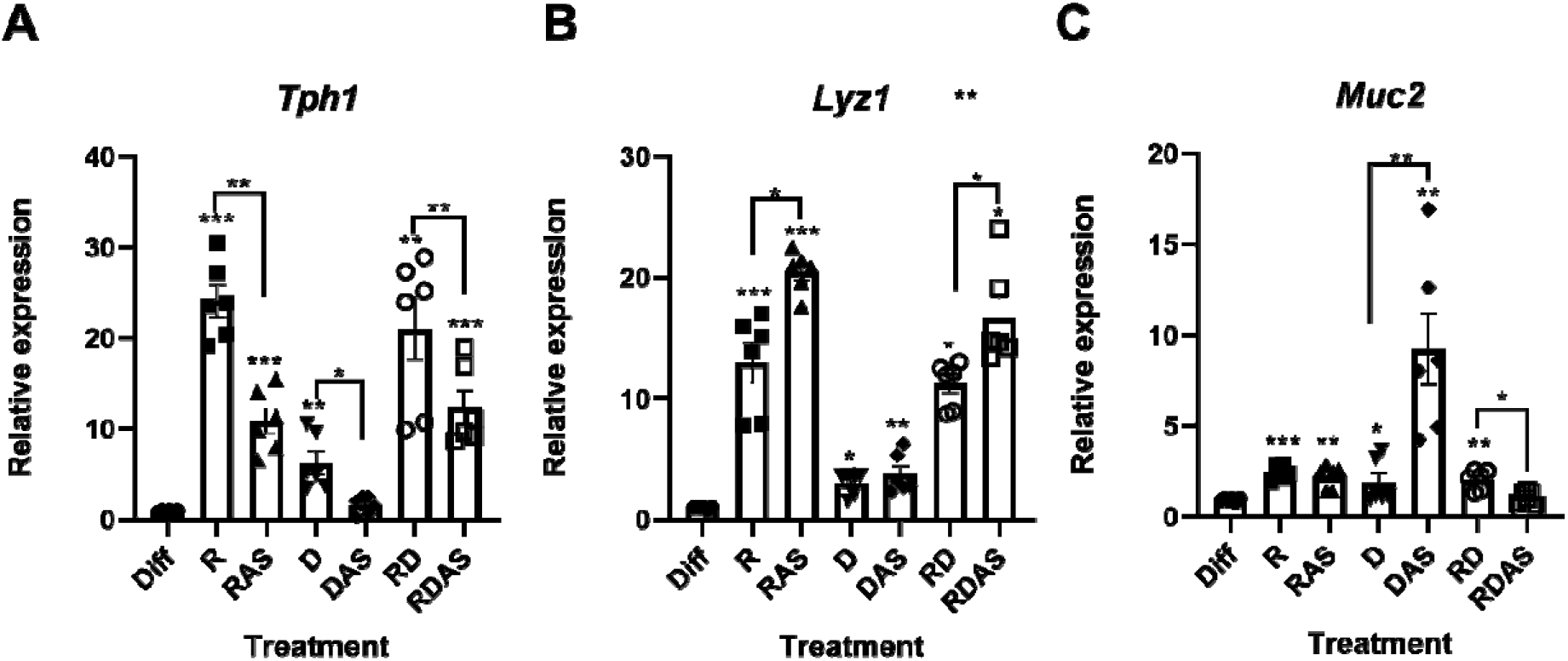
QPCR analysis showing *Tph1, Lyz1* and *Muc2* expression in mGO following treatment with the differentiation cocktail (n=6, bar graphs represent means ±SEM, paired t-test, * P<0.05; ** P<0.01; *** P<0.001).

## Lead Contact and Materials Availability

Further information and requests for resources and reagents should be directed to and will be fulfilled by the Lead Contact, Domenico Accili.

## Methods

### Animal studies

Tg(Neurog3-cre)C1Able/J (#005667, Ngn3-Cre)(Schonhoff *et al.*, 2004) and B6.Cg-Gt(ROSA)26^Sortm9(CAG-tdTomato)Hze^/ (#007909) were from the Jackson Laboratories. Tg(Ins2-cre)23Herr (Kuo et al., 2019) and FoxO1^fl/fl^ mice (Matsumoto et al., 2007) have been described. Tph1^CreERT2^ mice were kindly gifted by Prof. Juanita L. Merchant of University of Arizona and Lyz1^CreER^ mice by Prof. Na Gao of Rutgers University (Yu et al., 2018). Mice were housed in 12h light/dark cycle (7AM/7PM) barrier facility with free access to water and food. All animal studies were approved by and overseen by Columbia University Institutional Animal Care and Use Committee (IACUC).

### Physiological Studies

A single injection of Streptozotocin (STZ, 170 mg/kg, S0130, Sigma, St. Louis, MO) was administrated intraperitoneally to induce diabetes in 6 to 8-week old male NFKO and littermate male FoxO1^f/f^ (WT) mice followed by High-Dose STZ Induction Protocol from Animal Models of Diabetic Complications Consortium (AMDCC) (http://www.AMDCC.org). Ad libitum blood glucose levels and body weight were monitored every 2-3 days after STZ injection. Tail blood glucose after 4h of fasting was measured on days 0 and 7 using Contour Next glucometer. Mice that were not hyperglycemic (Fast Blood Glucose > 250 mg/dl) 7 days after STZ administration were excluded from further study. Mice were randomized base on blood glucose and body weight.

For in vivo drug treatment, mice were injected intraperitoneally with 25mg/kg DBZ once per day for 2 days or gavaged with 10mg/kg Repsox once per day for 5 to 7 days. Weight and ad libitum glucose levels were monitored every 2 days during treatment. In the fasting-refeeding study, mice were fasted for 4 hours followed by 1 hour refeed. Blood was collected from tail vein for glucose and insulin measurement. Plasma insulin was measured by enzyme-linked immunosorbent assay (ELISA, Mercodia, Uppsala, Sweden). In oral glucose tolerant test (OGTT), mice were fasted for 4 hours followed by gavaging of 2 g/kg (10 ml/kg) of D-glucose (Sigma) dissolved in distilled water. Glucose was measured at 0, 15, 30, 60 and 120 min by tail vein sampling.

### Human Tissues

Surgically resected intestinal tissues or endoscopic biopsy samples were obtained from 8 patients from the Columbia University Irving Medical Center and Presbyterian Hospital and Vanderbilt Clinic. The institutional review boards at Columbia University have approved all procedures and all samples were obtained with informed consent. All human fetal tissue samples were collected under Institutional Review Board Approval at both the University of Southern California and Children’s Hospital Los Angeles. Consent for tissue donation was obtained after the patient had already made the decision for pregnancy termination by Dilation and Curettage or Dilation and Evacuation and was obtained by a different clinical staff member than the physician performing the procedure. All tissue was de-identified, and the only clinical information collected was gestational age and the presence of any material of fetal diagnoses. Intestinal samples ranging in age from 15 to 17 weeks of gestation were received immediately after elective terminations and fixed in 4% paraformaldehyde, dehydrated with 30% sucrose, processed for OCT embedding, followed by sectioning and immunostaining.

### Chemicals

All small molecule information for intestinal treatment is listed in Table S1. All drugs were dissolved in Dimethylsulfoxide (DMSO) and diluted into previously indicated concentration with medium during the treatment. For in vivo treatment, dibenzazepine (DBZ) was from Apexbio Technology (Houston, TX); RepSox from Selleck Chemical (Houston, TX). DBZ and Repsox were formulated in 1% DMSO, 0.5% methylcellulose and 0.2% Tween-80 PBS solution, respectively.

### Gut Organoid Cultures

For mouse small intestinal organoids, crypts were isolated using EDTA chelation from the duodenum as described (Sato and Clevers, 2013) and cultured in complete growth medium that included advanced Dulbecco’s modified Eagle’s medium and F12, DMEM/F12 (Gibco, Grand Island, NY) containing penicillin or streptomycin 100 U/mL, HEPES 10 mmol/L, Glutamax 2 mmol/L, B27 supplements (Gibico), murine epidermal growth factor (mEGF) 50 ng/mL (Peprotech, East Windsor, NJ), N-acetylcysteine 1 mmol/L (Sigma), R-spondin-1 (conditioned medium 1ml/50ml, gifted by Prof. Calvin J. Kuo, Stanford University, CA), Noggin (conditioned medium 1ml/50ml, gifted by Prof. Hans Clevers, Hubrecht Institute, Netherlands). Differentiation medium consisted of the same components as complete growth medium without R-spondin-1, Noggin and hEGF.

For human small intestinal organoids, crypts were isolated using EDTA chelation from the duodenum as previously described (Fujii et al., 2015). IntestiCult™ Organoid Growth Medium (Human) or IntestiCult™ Organoid Differentiation Medium (Human) were used for culture or differentiation of human gut organoids (STEMCELL Technologies, Vancouver, Canada). Organoids were used prior to passage 3 for optimal efficiency of EEC and β-like cell induction.

### Intestinal epithelial cell isolation and sorting

4- to 6-week-old NFKO mice were used to isolate single intestinal cell preparations as described (Magness et al., 2013). A segment of the proximal intestine representing a 5-cm region from 2 to 7 cm distal to the pyloric sphincter was used for cell isolation. Attached pancreata were carefully removed under a dissection microscope to avoid pancreatic β-cell contamination. Isolated intestinal epithelial cells (10^7^ cells/1ml) were incubated with Mouse BD Fc Block™ (55314, BD, Franklin Lakes, NJ) for 10 minutes on ice in IESC media (DMEM/F12, B27, Glutamax, Penicillin/Streptomycin, 500 mM N-acetyl-cysteine and 10μM Y27632). Cells were stained 20 min with APC-conjugated anti-CD24 antibody (1:100 dilution, 138506, Biolegend, San Diego, CA) and FITC-conjugated anti-Epcam antibody (1:500 dilution, 118208, Biolegend), washed twice with IESC media and DAPI (0.1 μg/ml, D1306, Invitrogen, Waltham, MA) was added prior to sorting using BD Influx.

### Flow cytometric analysis of epithelial cells

Single cell suspension was obtained by enzymatic digestion of intestinal mucosa or cultured organoids (Magness *et al.*, 2013; Sato et al., 2011a). Suspended cells were first stained with live/dead cell staining kit (Invitrogen, L34964), then fixed in BD Cytofix™ fixation buffer (BD, 554655) for 20 min. Cells were washed in permeabilization buffer (0.2% Saponin, 0.5% BSA and 2mM EDTA in PBS) twice followed by intracellular primary and secondary antibody staining. After 3 washes with permeabilization buffer, cells were resuspended in FACS buffer (0.5%BSA and 2mMEDTA in PBS) for sorting or FACS analysis. When sorted cells were used for RNA isolation, 0.2% RNaseOUT (Invitrogen, 10777-019) was added to the antibody incubation and FACS buffer before sorting.

### RNA isolation and quantitative PCR analysis

Cultured gut organoids or FACS sorted cell were in 1ml TRIzol (ThermoFisher, Waltham, MA). RNA was isolated using RNeasy mini kit or RNeasy Micro kit (Qiagen, Germantown, MD) followed by reverse transcription using qScript cDNA Synthesis Kit (QuantaBio, Beverly, MA). RNA isolation from intercellular stained cell sample was as described (Thomsen et al., 2016). QPCR was performed with GoTaq® qPCR Master Mix (Promega, Madison, WI). QPCR primers sequence will be provided upon request. Gene expression levels were normalized to HPRT (hypoxanthine phosphoribosyl transferase) using the 2^-ΔΔCt^ method and are presented as relative transcript levels.

### Quantitative measurement of conversion insulin-producing cells with cultured organoids

Primary gut crypts which derived from a mouse bearing an Rip-Cre;Rosa26^tdTomato^ reporter allele placed in culture, and then induced to undergo cell conversion by applying a protocol based on published patent US20170349884A1. Briefly, the first 2 days after seeding the organoids, crypts cultured in complete growth medium with CHIR99021, Valproic acid and Y-27632 to enhance the stem cell vitality. At Day3, DAPT, Repsox, ISX-9 and Adc were added to complete growth medium. At Day 4-5, organoids were treated with differentiation medium supplemented with DAPT, Repsox, ISX-9, Adc, and IWP-2 to induce cell conversion. At Day 6, organoids were incubated in differentiation medium with DAPT, Repsox, PD0325901, T3, CHIR 99021 to induce further maturation of β-like cells. During the differentiation process (Day 4-6), a FOXO1 inhibitor (e.g., AS or one of the FBT compounds) was added to the medium along with the inhibitor cocktails listed above. Thereafter, cells were analyzed by flow cytometry, and cells in which *Ins2* expression has been activated were sorted from other epithelial cells by flow cytometry. These cells were collected by flow, RNA extracted and analyzed for expression of β-cell-specific genes. To evaluate the efficiency of cell conversion to INS2-positive cells we calculated a Tom Score by multiplying INS2 tomato intensity, percentage of INS2 Tomato cells, and percentage of live cells.

### In Situ Hybridization by RNAscope

RNAscope was performed using the RNAscope® 2.5 HD Detection Reagent – RED kit (Advanced Cell Diagnostics, Hayward, California) combined with immunofluorescence according to manufacturer’s instructions. A human insulin probe (ACD catalogue number 313571, NCBI reference sequence NM_000207.2) was used to detected insulin mRNA.

### Immunohistochemistry

Small intestine was isolated from 6 to 8-week-old or drug-treated mice. After harvesting, the tissues were rinsed with PBS, coiled into Swiss rolls, and fixed in 4% PFA for 2 hours, followed by dehydration in 30% sucrose in PBS overnight, embedding in Tissue-Tek O.C.T (Sakura, Torrance, CA), and freezing at – 80°C. Mouse and human organoids sections were prepared as described (Bouchi *et al.*, 2014). 6μm-thick sections were cut and stained with standard frozen-IHC protocols. The primary and secondary antibodies used are listed in Table S2. Images were recorded with a confocal laser-scanning microscope (LSM 710, Carl Zeiss, NY) and processed using Image J software (National Institutes of Health).

### Bulk RNA-sequencing and analysis

RNA-sequencing was performed by the Columbia Genome Center. Poly-A pull-down was used to enrich mRNA from small intestinal epithelial cells sorted from 4 to 6-week-old NFKO or Ins2-tomato^+^ cells from drug-treated mice. Library construction was done using Illumina TruSeq chemistry (intestinal epithelial cells) or the Clontech Ultra Low v4 kit for cDNA synthesis followed by NexteraXT (sorted cells from organoids). Libraries were sequenced using Illumina NovaSeq 6000. Samples were multiplexed in each lane, yielding targeted numbers of paired-end 100bp reads for each sample. RTA (Illumina) was used for base calling and bcl2fastq2 (version 2.19) for converting BCL to fastq format, coupled with adaptor trimming. A pseudoalignment to a kallisto index was created from transcriptomes (Mouse: GRCm38) using kallisto (0.44.0). Differentially expressed genes were tested under various conditions using Sleuth or DESeq2, R packages designed to test differential expression between two experimental groups from RNA-seq counts data. Pathway enrichment was assessed through the pre-ranked version of Gene Set Enrichment Analysis (GSEA) (Subramanian et al., 2005) and through Gene Ontology Analysis using David 6.8 Software (GO) (Huang et al., 2009).

### ScRNA-seq and data analysis

Tomato+ cells were isolated and sorted from NFKO mice as described above. Viability >90% was obtained for all samples as determined by trypan blue staining. 10x Genomics 3’ Single Cell Gene Expression (GEX) microfluidic cell processing, library preparation and sequencing were performed by the Columbia Genome Center as described (Capdevila et al., 2020). The raw counts of scRNA-seq data were analyzed as follow. The R package Seurat was used to do the clustering analysis and cell type annotation. (Satija et al., 2015). For the quality control, cells with either the percentage of reads mapped to the mitochondrial genome higher than 15%, or total UMI counts higher than 1e-6, or number of feature RNA smaller than 200 were removed. The “LogNormalize” normalization method was used to normalize library size. Clustering analysis was done using the unsupervised graph-based function of the Seurat package. First, the highly variable genes were identified, and data scaled. Then principal component analysis (PCA) was performed on the scaled data using highly variable genes. The neighbors of cells were found based on the first 50 PCs. The modularity was optimized by the Louvain algorithm. Finally, the clustering results are visualized on PCA plot. The integration method in Seurat was used for annotating the cell types based on known cell types in reference dataset (Stuart et al., 2019). Single-cell transcriptome data of mouse small intestinal epithelium was obtained from GSE92332 and used as reference (Haber et al., 2017). The differentiation potential of INS+ and INS_neg_ cells from NFKO mice was predicted using CytoTRACE (Gulati et al., 2020).

### Statistical analysis

Data analysis was conducted using Prism 6.0 software (GraphPad Software, San Diego, California) unless otherwise stated. Sample sizes were determined by the reproducibility of the experiments and are similar to those generally employed in the field. Statistical significance in two-way comparisons was determined by a Student’s t-test or paired t-test when data met the normal distribution, whereas ANOVA was used when comparing more than two datasets. If the data did not meet the normal distribution, the Wilcoxon test or Mann-Whitney test were used. The statistical test and significance level are indicated in the figure legends. Asterisks denote statistically significant differences from the control. (*) p < 0.05; (**) p < 0.01; and (***) p < 0.001.

